# Active sampling state dynamically enhances olfactory bulb odor representation

**DOI:** 10.1101/222588

**Authors:** Rebecca Jordan, Izumi Fukunaga, Mihaly Kollo, Andreas T. Schaefer

## Abstract

The olfactory bulb (OB) is the very first site of odor information processing, yet a wealth of contextual and learned information has been described in its activity. To investigate the mechanistic basis of contextual modulation, we use whole-cell recordings to measure odor responses across rapid (<30 min) learning episodes in identified mitral/tufted cells (MTCs). Across these learning episodes, we found that diverse response changes occur already during the first sniff cycle. Motivated mice develop active sniffing strategies across learning, and it is this change of active sampling state that dynamically modulates odor responses, resulting in enhanced discriminability and detectability of odor representation with learning. Evoking fast sniffing in different behavioral states demonstrates that response changes during active sampling exceed those predicted from purely feed-forward input. Finally, response changes are highly correlated in tufted cells, but not mitral cells, indicating cell-type specificity in the effect of active sampling, and resulting in increased odor detectability in the tufted and enhanced discriminability in the mitral cell population over the rapid learning episodes. Altogether, we show that active sampling state is a crucial component in modulating and enhancing olfactory bulb responsiveness on rapid timescales.

## Introduction

The ability to respond to sensory stimuli according to learning and context is vital for orchestrating appropriate behavior. Our view of sensory processing has shifted away from the simplicity of passive feed-forward models driven by sensory stimuli, to one that additionally incorporates contextual information provided by top-down circuits into the ongoing processing (Engel et al., 2001). This has been driven in part by observations that activity in primary sensory cortex is widely modulated by contextual information: locomotion, attention, reward timing and experience all modulate visual cortex activity (Chubykin et al., 2013; Fiser et al., 2016; Ito and Gilbert, 1999; Niell and Stryker, 2010), while whisking behavior and social context modulate barrel cortex activity (Crochet and Petersen, 2006; Ferezou et al., 2006; Lenschow and Brecht, 2015).

The olfactory bulb (OB) is the very first site of odor information processing, yet already modulation of OB neural output by a wealth of contexts and behavioral tasks has been described from recordings of suprathreshold activity, including unit recordings, calcium imaging, and LFP recordings. These include modulation of odor responses by hunger state (Pager, 1974; Pager et al., 1972), task-engagement (Fuentes et al., 2008), reward anticipation (Doucette and Restrepo, 2008), conditioned aversion (Kass et al., 2013), and even non-olfactory events (Kay and Laurent, 1999; Rinberg et al., 2006). Recently, a number of studies have described changes in mitral and tufted cell (MTC) odor responses over the course of olfactory learning (Chu et al., 2016; Doucette and Restrepo, 2008; Yamada et al., 2017). Despite the prominence of such studies, the mechanistic basis underlying contextual modulation of the circuit is still unclear. In particular, rarely have these contextual modulations been interpreted in the framework of active sampling behavior, which is known to be controlled in a complex and context dependent manner (Wachowiak, 2011), including over the learning of olfactory tasks (Kepecs et al., 2007; Wesson et al., 2008, 2009; Youngentob et al., 1987). Not only this, but unit recordings as well as imaging do not have access to subthreshold activity, while the former also has the potential to misidentify cell types and bias recordings toward a subpopulation of MTCs that have high baseline firing rates (Kollo et al., 2014).

To investigate the mechanistic basis of task dependent changes in mitral and tufted cell odor responses, we recorded from identified mitral and tufted cells using blind whole-cell recordings *in vivo* in a range of behavioral states. We optimised training protocols for an olfactory task to facilitate very rapid olfactory discrimination learning episodes, which allowed us to make whole-cell recordings over the full learning epoch. At the same time, we measured sniffing behavior using an external flow sensor. Altogether, we provide evidence that learned active sampling behavior overtly modulates olfactory responses in a cell-type specific way that cannot be explained by feed-forward input, and overall appears to enhance the representation of odors across the olfactory bulb.

## Results

### There are differences in odor responses according to behavioral state

We recorded from 23 MTCs in passive mice exposed to repeated stimulation of odor mixtures (Figure S1), as well as 21 MTCs in mice during learning of a simple olfactory go/no-go discrimination task with the same mixtures. In our task-learning mice, after pre-training on different odorants (Figure S2A), mice underwent very rapid learning on a novel pair of odor stimuli, reaching criterion within 10-20 minutes (Figure S2B). It was thus possible to make stable whole-cell recordings over the full timescale of learning. MTCs were distinguished from interneurons as previously described (Kollo et al., 2014), using independent component analysis of the AHP waveform. This was confirmed with morphological reconstruction of 9 MTCs (Figure S3; see methods).

To first determine whether the two behavioral states cause any general change in olfactory bulb physiology, we applied a series of current steps and compared the basic properties of cells between the passive and learning states. Both the passive properties (resting membrane potentials, input resistance and membrane time constants) and spontaneous activity (spontaneous firing rates and sniff phase-modulation of membrane potential – ‘sniff-V_m_ mod.’) of cells revealed little detectable difference in either average values or variance (see supplemental information; Figure S4A-F).

Next, basic odor response properties were compared between MTCs in passive (Figure 1A; 46 cell-odor pairs) and task-learning mice (Figure 1B; 42 cell-odor pairs). Note that all odor responses in the manuscript are aligned on each trial to the first inhalation onset. Comparing passive and learning cohorts by averaging responses across all trials for a given cell-odor pair revealed that firing rate (FR) responses did not overtly differ in distribution between passive and behaving mice (Figure S4G-H). Median FR responses were similar (passive: median = −0.84 Hz, IQR = −2.2-1.1 Hz; learning: median = −0.51 Hz, IQR = −2.8-2.1 Hz, p = 0.84, Ranksum), as was variance across cell-odor pairs (p = 0.42, Brown-Forsythe test). Measuring a cell’s input using subthreshold responses offers us a more sensitive measure of many response parameters, including temporal features and inhibition. Taking average membrane potential responses revealed that these do not differ much between the two behavioral states in terms of means (passive: mean = −1.5 mV, SD = 1.8 mV; learning: mean = −1.7 mV, SD = 2.4 mV; p = 0.67, unpaired t-test; Figure 1C_i-ii_), however response variance was larger across cells in learning mice relative to passive mice, due to higher representation of both strong inhibition and excitatory responses (p = 0.05, unpaired t-test, n = 42 vs 46; Figure 1C_ii_).

**Figure 1.**
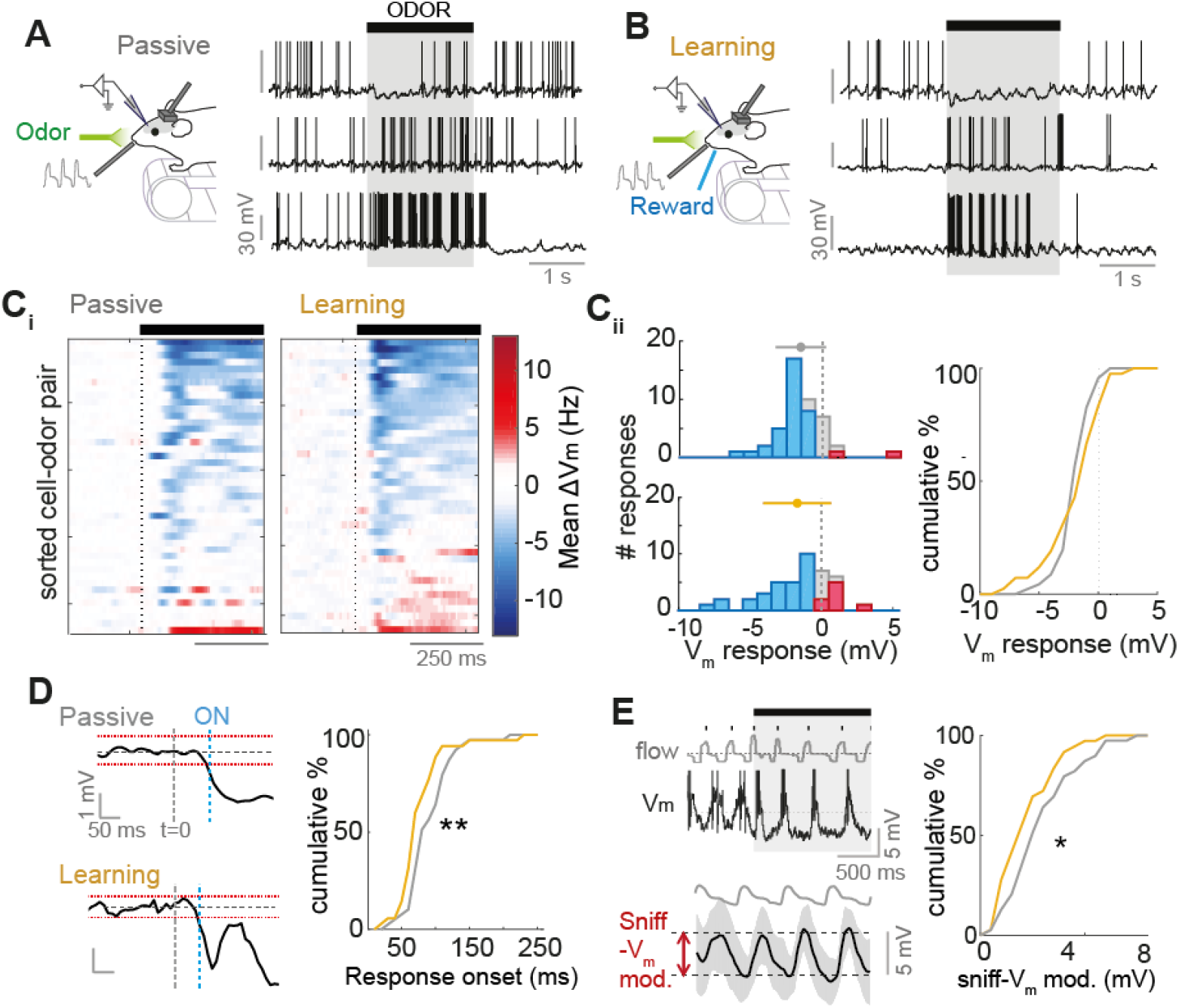
Differences in odor responses according to behavioral state. (**A**) Example odor response traces for three different cell-odor pairs recorded in passive mice, aligned to first inhalation onset. (**B**) As for panel A, but for cell-odor pairs recorded in learning mice. **C**_i_) Heatmap of V_m_ responses averaged across all trials for each cell-odor pair, sorted by mean 500 ms V_m_ response, for both passive (n=46) and learning (n=42) datasets. Black bar indicates odor stimulus, aligned to first inhalation onset. (**C**_ii_) Left: Histograms of average 500 ms V_m_ responses for (top) passively exposed and (bottom) learning mice. Right: cumulative histograms comparing average V_m_ response data for passive (grey) and learning (gold) cell-odor pairs. (**D**) Comparison of response onset latency for learning and passive mice. Left shows examples: average membrane potential waveforms averaged over all trials. t=0 indicates when the odor turned on, aligned to the first inhalation onset. Red dotted lines indicate upper bounds and lower bounds (calculated as mean ± SD of the baseline membrane potential). When the V_m_ waveform rises above or below the upper or lower bound respectively for at least 50 ms, this is when response onset is defined (blue dotted line, ON). Right: cumulative histograms to compare response onsets for passive and learning mice. (**E**) Left: example of a highly sniff-locked odor response from a passive mouse across the first 4 sniff cycles. ‘Flow’ shows nasal flow trace. Below trace shows example average phase aligned membrane potential for first four sniffs of the odor response. Shaded area shows SD. ‘Sniff-V_m_ mod.’ indicates the calculation of sniff-V_m_ modulation amplitude (see methods). Right: Cumulative histograms to compare sniff-modulation amplitudes for passive and learning mice.

We next compared temporal features of the subthreshold responses. Comparing response onsets between passive and learning mice revealed a significant shift towards earlier onsets in learning mice (passive: median = 85 ms, IQR = 70-110 ms, n = 39; learning: median = 70 ms, IQR = 60-90, n = 36; p = 0.004, Ranksum; Figure 1D), with 33% of responses occurring before 70 ms in learning mice, and only 10% in passive mice. Just as for baseline activity (Figure S4F), activity during odor response is often locked to the sniff cycle. We calculated the amplitude of membrane potential modulation when aligned to sniff phase (sniff-V_m_ modulation amplitudes) during the odor response (Figure 1E; see methods for details) to quantify to what degree each cell-odor pair was locked to the sniff cycle during odor stimulation. Overall, passive cell-odor pairs showed a significantly higher degree of patterning by the sniff cycle than learning cell-odor pairs (passive sniff-V_m_ modulation amplitude=3.1±1.7 mV, n=42; learning: 2.4±1.4 mV, n=38, p=0.03, unpaired t-test; Figure 1E).

Overall, this analysis revealed that the most overt differences between passive and learning mice were measurable in subthreshold responses, which showed increased variance, shorter latency and less sniff coupling in learning mice. We thus focused primarily on subthreshold responses for the next set of analyses.

### Diverse odor response changes occur within the very first sniff cycle in learning mice

Recent imaging work has suggested that MTC responses are subject to change in both learning and passive mice over long timescales (Chu et al., 2016; Doucette and Restrepo, 2008; Yamada et al., 2017). To assess whether the increased response variance apparent in learning animals (Figure 1Cii) developed across rapid go/no-go task learning (Figure 2A), we compared the subthreshold response of each cell-odor pair in early trials where the mouse is performing at chance levels, with the response in late trials where the mouse is performing at criterion or above (Figure 2B). Since median reaction times in the task were 500 ms (Figure S2C), we focused our analyses on the first 500 ms of odor response.

**Figure 2.**
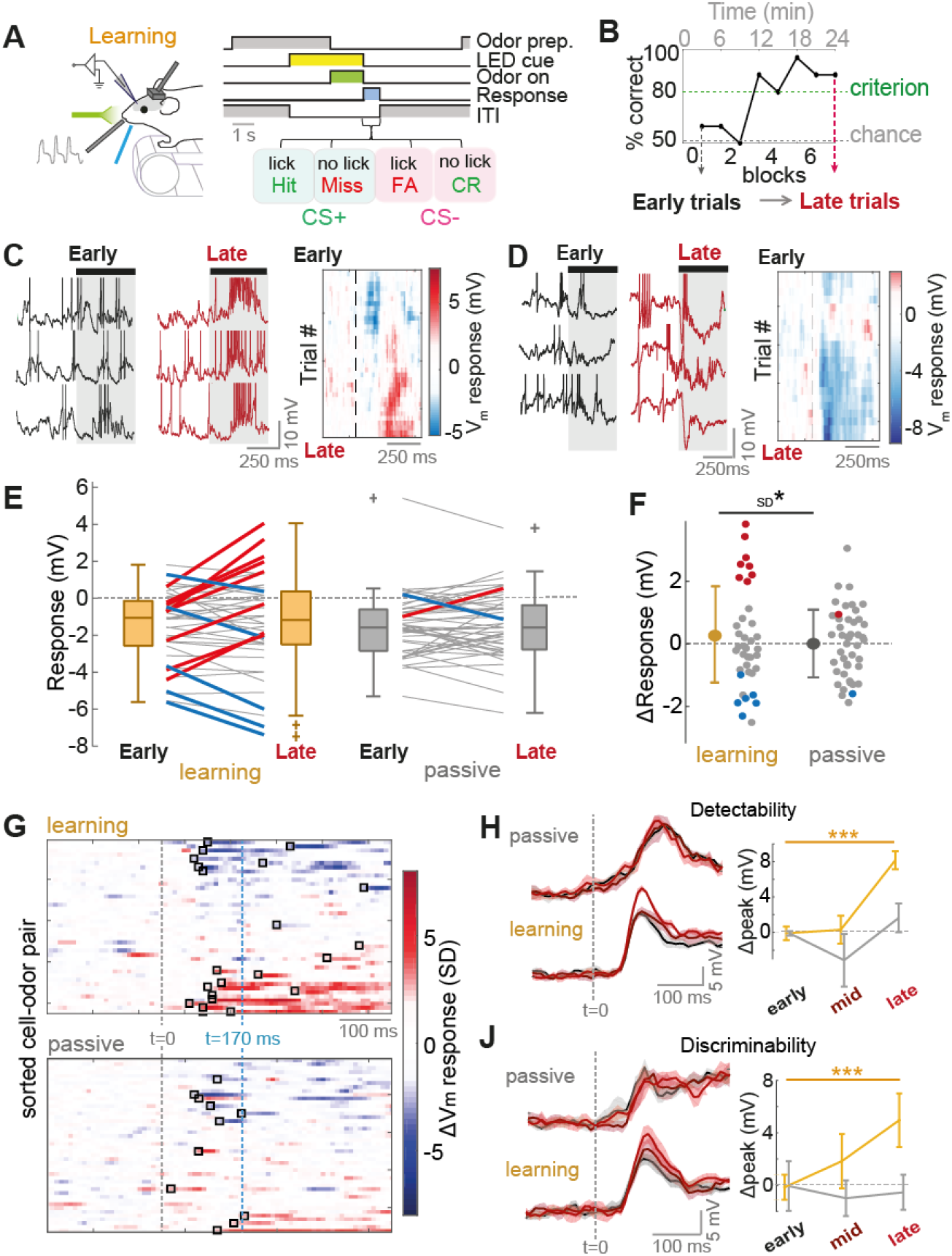
Diverse odor response changes occur within the very first sniff cycle in learning mice. (**A**) Diagram of the recording paradigm (left) and schematic of go/no-go task sequence (right). (**B**) Example learning curve for one mouse across the recording timeframe. Responses are compared between five early trials (unlearned) and five late trials (learned) to assess learning-related changes. (**C**) Left: example odor response traces in early and late for a cell-odor pair undergoing increase in excitation across learning (spikes have been clipped). Shaded area indicates odor stimulus (aligned to first inhalation onset). Right: heatmap showing 5-trial moving average of membrane potential response across trials (**D**) As for panel C, but for a response undergoing an increase in inhibition. (**E**) Plot of early and late membrane potential responses (first 500 ms) for learning mice (left; n=42 cell-odor pairs) and passive mice (right; n=46 cell-odor pairs) separately. Thick red lines indicate significant positive change (p<0.01), thick blue lines indicate significant negative change. (**F**) Comparison of response changes (late-early) for learning and passive mice. Red dots show significant positive changes, blue dots show significant negative changes. (**G**) Response change heatmaps (late-early average membrane potential waveforms) normalized by baseline SD. Black boxes indicate onset of change (>2 SD for at least 5 consecutive points). T=170 ms is indicated as the minimal detected reaction time. (**H**) Left plots: Euclidean distance as a function of time since odor onset (t=0, aligned to first inhalation onset) between population vectors for odor response, and control vectors initiated by an inhalation during the inter-trial interval. This gives an indication of the detectability of the odor response across the sample. Black plot is calculated from early trials, maroon plot is calculated from mid-point trials, and red plot is calculated from late trials. Each is averaged over 5 trial subsets (see methods for details), and shaded area indicates standard deviation. Right: plot to show peak detectability within the first 170 ms of the stimulus across early, mid-point and late trials. Plot shows mean across the 5 trial subsets, and errorbars show standard deviation. Gold plot is for learning mice (n=42 cell-odor pairs), and grey plot is for passive mice (n=46 cell-odor pairs). (**J**) As for H, but with the Euclidean distance measured between population vectors for the CS+ and CS− to give an indication of the discriminability of the two responses across the sample.

We noticed that there were diverse changes in odor response occurring over the course of learning: for example, overt increases in excitatory response (Figure 2C and S5A), as well as increases in inhibitory response (Figure 2D), which developed across trials. Overall, in learning mice, 30% (13/42) of cell-odor pairs showed a significant change across learning (p<0.01, unpaired t-test between 5 early and 5 late trials), with 19% (8/42) showing a positive change, and 11% (5/42) showing a negative change (Figure 2E; Figure S5C). These changes led to an increase in the diversity of responses between early and late trials across the sample, though this did not quite reach significance (early SD = 2 mV, late SD = 2.6 mV; p = 0.06, Bartlett test). On a trial by trial basis, changes in excitatory subthreshold response were reflected by changes in firing rate, though this was not clear for changes in inhibition (Figure S5A-B). In contrast to learning mice, cell-odor pairs recorded in passive mice showed far less frequent significant changes (4%, 2/46 cell-odor pairs), and no change in variance across the sample (SD early = 1.9 mV, SD late = 2.0 mV, p = 0.58, Bartlett test; Figure 2E and S5D). Overall, there was significantly higher variance in response changes for cell-odor pairs recorded in learning compared to passive mice (learning ΔV_m_ SD = 1.5 mV; passive ΔV_m_ SD = 1.1 mV; p = 0.02, Bartlett test; Figure 2F). Learning-related changes were not due to time-dependent effects of recording, since recording durations in passive and learning were matched (Figure S5E). Response changes did not reflect the contingency of the odor or response of the animal, since the distribution of changes showed no significant difference between CS+ and CS− stimuli (p = 0.77, paired t-test; Figure S5F), and were even correlated (R^2^ = 0.44, p = 0.001; Figure S5G).

What aspects of the response change could potentially be used to aid decision making? Mice are known to make simple olfactory discriminations within the timescale of a single sniff cycle. Congruently here, we find reaction times as low as 170 ms (Figure S2C-D). By identifying the onset of response change (see methods), we could show that 71% of identifiable changes occurred prior to 170 ms (median ΔV_m_ onset = 120 ms, IQR = 20-220 ms, Figure 2G and S5H), and 45% occurred prior to the 1st percentile of sniff durations (107 ms; Figure S5J). Thus, changes occur within the timescale of a single sniff cycle, and therefore could contribute to decision making. To assess the functional consequence of the learning-related changes for odor representation within this short timescale, we constructed a population response vector from the full sample of cell-odor pairs (similar to Figure S5C-D) and calculated the Euclidean distance of this population response vector from baseline data (see methods for details). We found that peak detectability within the first 170 ms of odor response significantly increased between early and late trials (peak early = 31.9 ± 0.8 mV, late = 40.1 ± 1.0 mV; p = 4×10^−5^, unpaired t-test, n-5; Figure 2H), while no such significant changes were observed for passive exposure (peak early = 35.3 ± 0.3 mV, late = 37.1 ± 1.6 mV; p = 0.06 unpaired t-test, n-5; Figure 2H). By calculating Euclidean distances between response vectors for CS+ and CS− stimuli, we also observed a significant increase in discriminability of the two odors across the recording for learning (peak early = 19.8 ± 1 mV, late = 25.0 ± 2 mV, p = 5×10^−4^; Figure 2J) but not passive mice (peak early = 20.4 ± 1.9 mV, late = 19.9 ± 1.3 mV, p = 0.68). Since both discriminability and detectability peaked within 100 ms from odor onset, this enhanced representation occurred within the timescale of the first sniff cycle.

Thus, diverse response changes specifically occur across learning occurring on the timescale of a single sniff cycle, giving rise to enhanced early odor representation.

### Active sampling strategies emerge across task learning

What are the mechanisms underlying these response changes? Odors are acquired from the environment through sniffing behavior, which is subject to complex contextual modulation (Kepecs et al., 2007; Wachowiak, 2011; Wesson et al., 2009). To analyze sniff changes within the short 500 ms time-window of the odor stimulus, we measured sniffing using an external flow sensor and quantified the mean inhalation duration (MID) of all inhalations completed within the first 500 ms of the stimulus (Figure 3A). When comparing early and late learning trials, we noticed that mice showed significant changes in sniff behavior during the odor stimulus, with faster, sharper inhalations emerging gradually across learning (reduced MID, Figure 3B). Reductions in MID mirrored increases in sniffing frequency across trials (Figure 3C), and are thus indicative of faster sniffing.

**Figure 3.**
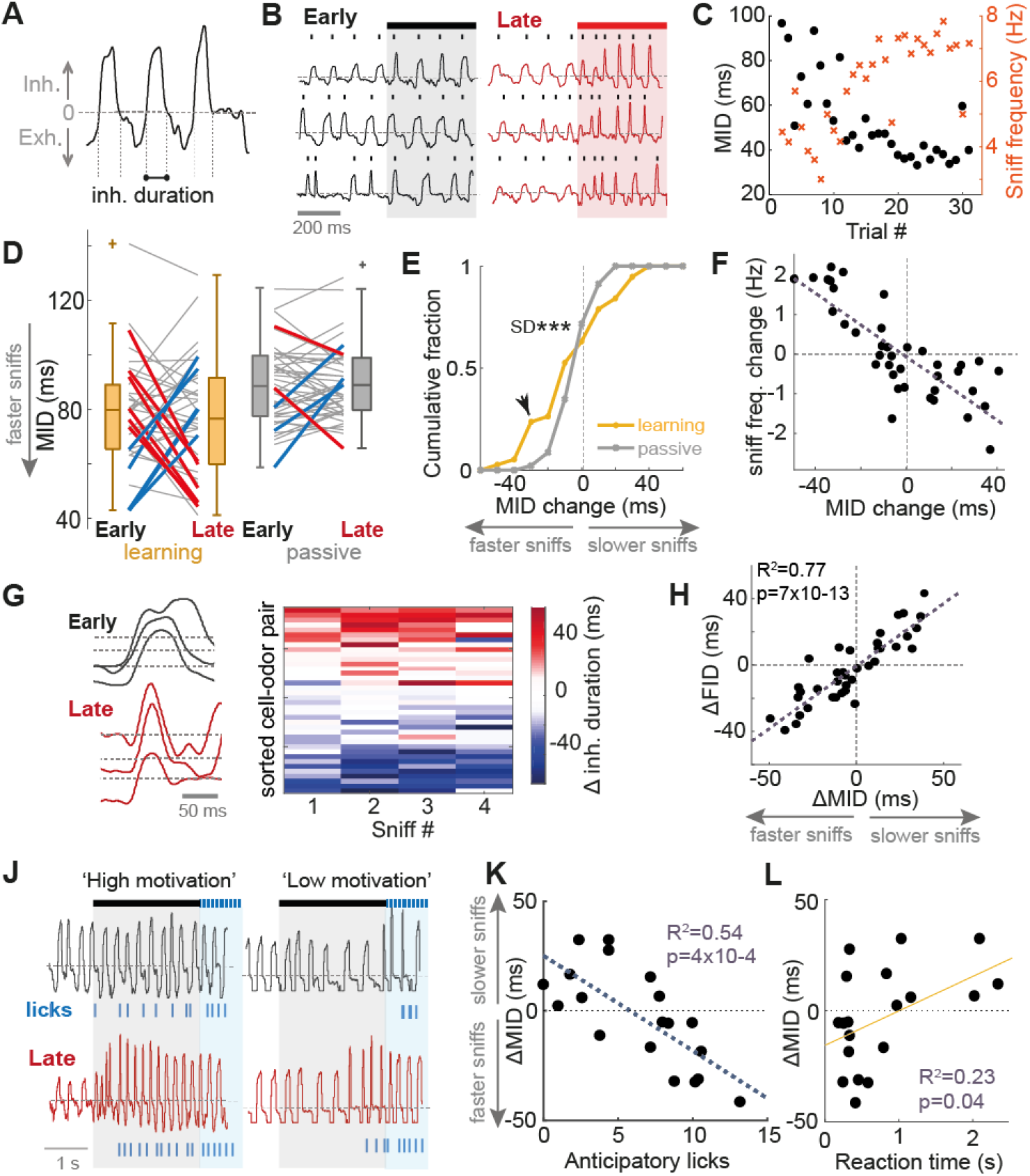
Active sampling strategies emerge across task learning. (**A**) Diagram to show extraction of inhalation duration from example nasal flow trace. (**B**) Example nasal flow traces from one mouse showing emergence of rapid sniffing between early and late trials. (**C**) MID for example cell-odor pair in panel B calculated for each trial (first 500 ms of stimulus) in black dots. Orange crosses show corresponding sniff frequency for each trial. (**D**) Plot showing how MID changes between early and late trials, for learning (n=38) and passive (n=42) mice. Thick red lines show significant reductions in MID (faster sniffing), thick blue lines show significant increases in MID (slower sniffing). (**E**) Cumulative histograms of MID change for learning and passive mice compared. Black arrowhead shows significant difference in the histograms (see methods). (**F**) Scatter between change in MID and change in sniff frequency between early and late trials for all learning cell-odor pairs. (**G**) Left: example flow traces showing duration of the first inhalation (FID) after odor onset between early and late trials. Right: heatmap to show change in inhalation duration as a function of sniff number since odor onset, sorted by change in MID. (**H**) Scatter of changes in FID versus changes in MID. (**J**) Example nasal flow traces during CS+ presentations for ‘high motivation’ (left) and ‘low motivation’ mice (right). ‘Motivation’ here refers to the number of licks during the odor stimulus (‘anticipatory’ licks). Note sniff changes only occur for the ‘high motivation’ mouse. (**K**) MID change (averaged for each cell across CS+ and CS− stimuli) across learning as a function of the mean number of anticipatory licks in late trials for CS+ trials. (**L**) MID change (averaged for each cell across CS+ and CS− stimuli) across learning as a function of the reaction time calculated from divergent lick patterns.

Across all cell-odor pairs, a large fraction underwent significant changes in MID during learning (26%, 10/38 cell odor pairs; Figure 3D). In stark contrast, passively exposed mice showed far more stable MID, with only 12% of cell odor pairs showing significant change (Figure 3D), and substantially less variation in the ΔMID; learning: SD = 24 ms, passive: SD = 9 ms, p = 3×10^−8^, Bartlett test; Figure S6A). Comparing cumulative histograms of the MID change between learning and passive mice revealed that a significantly larger proportion of learning mice underwent reductions in MID exceeding 20 ms (learning: 26%, passive: 2%; p<0.01, bootstrapping; Figure 3E). Changes in MID across the population of learning mice again correlated very well with changes in sniff frequency (R^2^ = 0.59, p = 3×10^−8^; Figure 3F).

In learning mice, changes in MID were highly correlated between rewarded and unrewarded odors (R^2^ = 0.54, p = 3×10^−4^, Figure S6B) and already the first inhalation after odor onset showed a pronounced reduction in duration (Figure 3G), with a tight correlation between changes in MID and changes in the first inhalation duration (FID; R^2^ = 0.77, p = 7×10^−13^; Figure 3H). Together this suggests that rapid sniffing in learning mice is likely to reflect an active sampling strategy rather than changes concomitant with reward anticipation or licking response.

What causes the variance in sniff changes across mice? Response vigour has previously been used as a measure of motivation levels in mice (Berditchevskaia et al., 2016). We noted that some mice would respond more eagerly to the CS+ stimulus than others, with larger frequency of anticipatory licking (licking 500-2000 ms after odor onset) in the late trials after learning was complete, while others would wait during the odor stimulus and only lick during the subsequent response period (Figure 3J). The number of anticipatory licks in late trials correlated well with the change in MID across learning, with reductions in MID associated with higher frequency anticipatory licking (R^2^ = 0.54, p = 4×10^−4^; MID change averaged across CS+ and CS− for each cell-odor pair; Figure 3K). Since the correlations existed for changes in MID for both CS+ and CS− alone (Figure S6C), these associations were not due to simple motor effects relating to the go response or reward expectation. Reduced MID was also significantly associated with shorter reaction time (Figure 3L; R^2^ = 0.23, p = 0.04, MID change averaged across CS+ and CS− for each cell-odor pair).

Thus, mice displayed changes in active sampling strategy forming across the learning session, with the development of fast sniffing associated with high motivation and short reaction times.

### Positive response changes are tightly linked to changes in active sampling

Since the MTCs recorded in awake animals were widely modulated by sniffing (Figure 1E), and mice displayed changes in sniff strategy (Figure 3), we next wanted to test what impact the changes in active sampling had on the response changes observed across learning. We first split the dataset according to MID change: large MID change (>20 ms absolute change between early and late trials, n=18), and small MID change (<20 ms absolute change, n=20). Comparing heatmaps of response change between early and late trials for each dataset revealed that positive changes were exclusively displayed alongside large MID change (Figure 4A). There was a significant increase in response variance for cell-odor pairs recorded alongside large MID change (early SD=1.8 mV, late SD=3.2 mV, p=0.02 Bartlett test), but not for small MID change (early SD=2.2 mV, late SD=2.2 mV; p=0.98 Bartlett test; Figure 4B). Altogether, responses recorded alongside large MID changes accounted for 7/8 significant positive response changes, and 2/5 inhibitory response changes, and showed significantly larger variance in response changes compared to those recorded alongside small MID changes (large sniff change: SD=1.9 mV, n=18; small sniff change: SD=1.1 mV, n=20; p=0.002, Bartlett test), and response changes in passive mice (passive SD=1.1 mV, p=0.002, Bartlett test, n=18 vs 46), while the small change dataset was indistinguishable from passive controls (p=0.94, Bartlett test, n=20 vs 46). In particular, there were significantly more positive response changes (>1 mV) in the large sniff change group (39%) compared to small sniff change (5%) and passive mice (11%; p<0.01, bootstrapping, see methods; Figure 4C).

**Figure 4.**
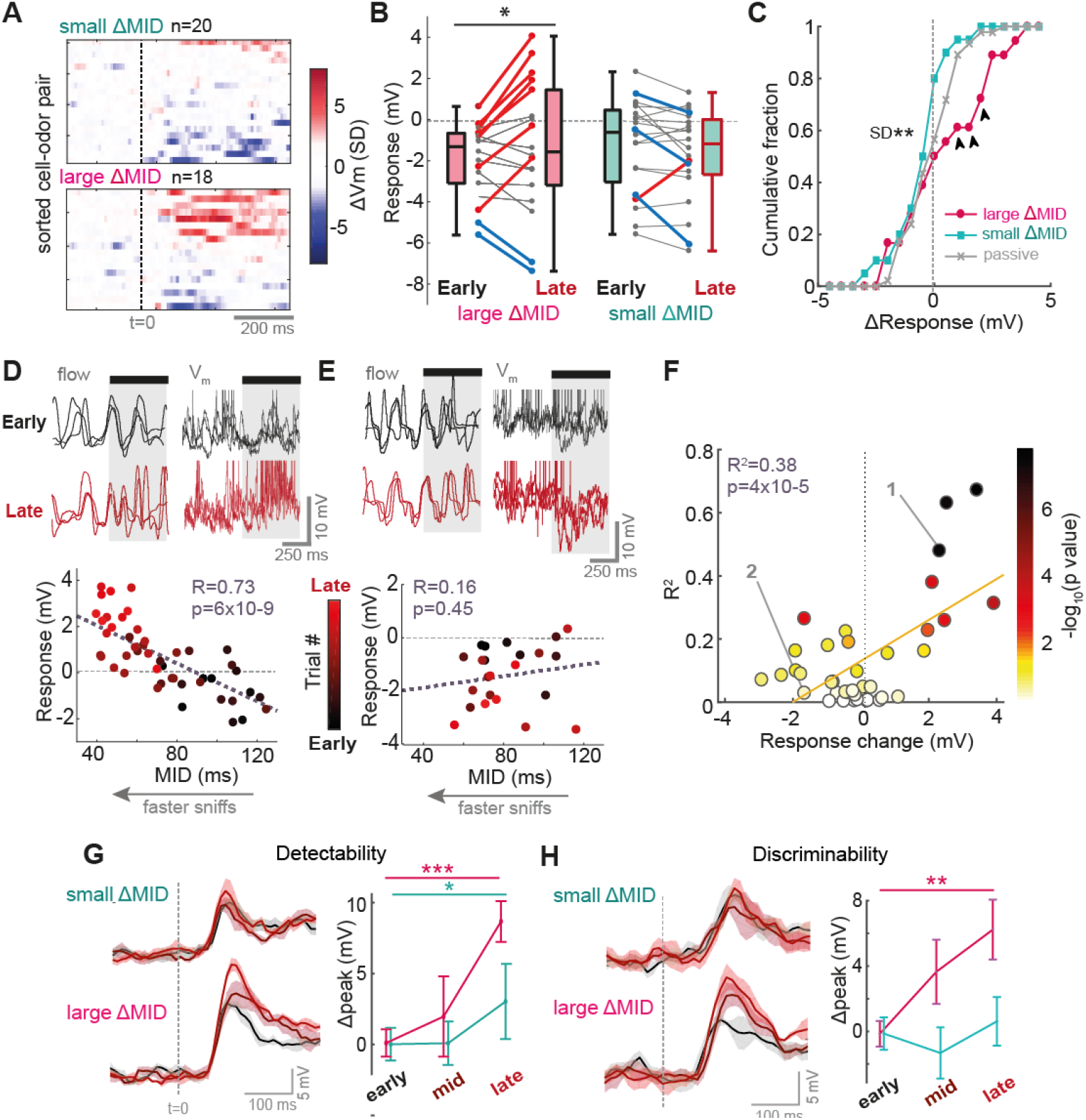
Positive response changes are tightly linked to changes in active sampling. All data is from the learning dataset. (**A**) Response change heatmaps (late-early average V_m_ response) normalized by baseline SD, for small MID change (ΔMID<20 ms) and large MID change (ΔMID>20 ms). (**B**) Plot of early and late membrane potential responses (first 500ms) across learning for large ΔMID (left; n=18 cell-odor pairs) and small ΔMID (right; n=20 cell-odor pairs) separately. Thick red lines indicate significant positive change (p<0.01), thick blue lines indicate significant negative change. (**C**) Cumulative histograms for response changes in large ΔMID, small ΔMID and passive mice. Black arrowheads show significant differences between large ΔMID vs both small ΔMID and passive histograms (see methods). (**D**) Above: example nasal flow and V_m_ traces overlayed for 3 early and 3 late trials for a cell-odor pair undergoing significant increase in excitation across learning. Spikes have been clipped for display. Below: Scatter between MID and V_m_ response across trials for this cell-odor pair. Points have been colored according to trial number. (**E**) As for panel D, but for a cell undergoing a significant increase in inhibition across learning. (**F**) Scatter between the response change across learning (late-early), and the R^2^ value for correlations as in panels D-E, colored according to the p-value of the correlation. Note how all cases of strong positive response change exclusively show strong correlations with sniffing. (**G**) Left: Euclidean distance between population response vectors and baseline data, now split into cell-odor pairs recorded alongside large MID (>20 ms change, n=18), and small MID (<20 ms change, n=20). Right: plot to show peak detectability within the first 170 ms of the stimulus across early, mid-point and late trials. Plot shows mean across the 5 trial subsets, and errorbars show standard deviation. (**H**) As for panel G, but for the Euclidean distance between population response vectors for CS+ and CS− for learning data that is now split into cell-odor pairs recorded alongside large MID change (>20 ms change for both CS+ and CS−, n=8 cells) and small MID change (any other cell, n=11 cells).

To test the strength of associations between V_m_ response and active sampling further, we correlated the mean MID and V_m_ response across trials for each cell-odor pair. For cells undergoing positive response changes across learning, this resulted in robust significant correlations, as in Figure 4D, while those undergoing increases in inhibition showed no such tight correlation (Figure 4E). Overall, 88% (7/8) cell-odor pairs showing significant positive changes across learning displayed highly significant correlations (p<0.01) between V_m_ response and MID across trials. Increased inhibition however could not be explained by active sampling changes, with no significant correlations between changes in V_m_ response and inhalation duration for these 5 cell-odor pairs. This effect across the population resulted in a significant positive relationship between the response change occurring across learning and the R^2^ of the correlation between MID and response across trials (R^2^=0.38, p=4×10^−5^, n=42; Figure 4F).

We also assessed whether sniffing could account for the differences in response onsets and sniff-V_m_ modulation amplitudes seen between passive and task-learning state (Figure 1D and E). Analysing these parameters over trials selected to match sniff parameters for each group demonstrated that differences in sniffing indeed accounted for differences in response onset and average sniff-V_m_ modulation amplitudes, although passive cell-odor pairs still showed a tendency toward very large sniff-V_m_ modulation amplitudes (Figure S7; see supplemental information).

How did changes in active sampling impact on changes in odor representation across the dataset? To test this we split the learning population according to MID change as before (Figure 4A-C). When recalculating the Euclidean distances for these individual datasets, we found that the increase in detectability largely occurred alongside large MID change (early peak=25.1±1.0 mV, late peak=33.7±1.4 mV; p=7×10^−4^, unpaired t-test; Figure 4G), and was far smaller and less significant in those undergoing small MID change (early peak=20.0±1.2 mV, late peak=23.1±2.6 mV; p=0.04, unpaired t-test; Figure 4G). We found the same result for changes in discriminability, with a significant increase only in cases where ΔMID for both CS+ and CS− stimuli was large (large ΔMID: peak early=13.2±0.8 mV, late=19.6±1.8 mV, p=0.002, unpaired t-test; small ΔMID: peak early=15.4±1.0 mV, late=16.1±1.5 mV, p=0.28, unpaired t-test; Figure 4H).

Thus, positive response changes are associated with changes in active sampling, which enhances early odor representation in terms of both detectability and discriminability while negative response changes (increased inhibition), cannot be explained by sniff changes.

### Active sampling and associated response changes are dynamically linked to task engagement

We next wanted to investigate the effect of dynamic changes in behavioral state on the changes in active sampling and odor responses observed. To do this, we recorded from 8 cell-odor pairs in an entirely new cohort of mice who were trained to criterion on the task prior to recording. If the rapid sniffing is indeed an active strategy for odor acquisition during behavior, we would expect the strategy to disappear if the task comes to an end (i.e. transition to passive odor exposure), and re-emerge when the task reinitiates. To test this, we implemented a paradigm in which task engagement could be reversibly changed by physically removing and re-introducing the water reward spout (Figure 5A), resulting in rapid switches between olfactory behavior and passive exposure as indicated by animal licking responses (Figure 5A and S8A-B). As predicted, animals robustly adapted their sniffing strategy upon elimination of the licking response after removal of the reward port (Figure 5B), with MID increasing (slower sniffing). Reintroduction of the reward port rapidly restored fast sniff behavior (reduced MID).

**Figure 5.**
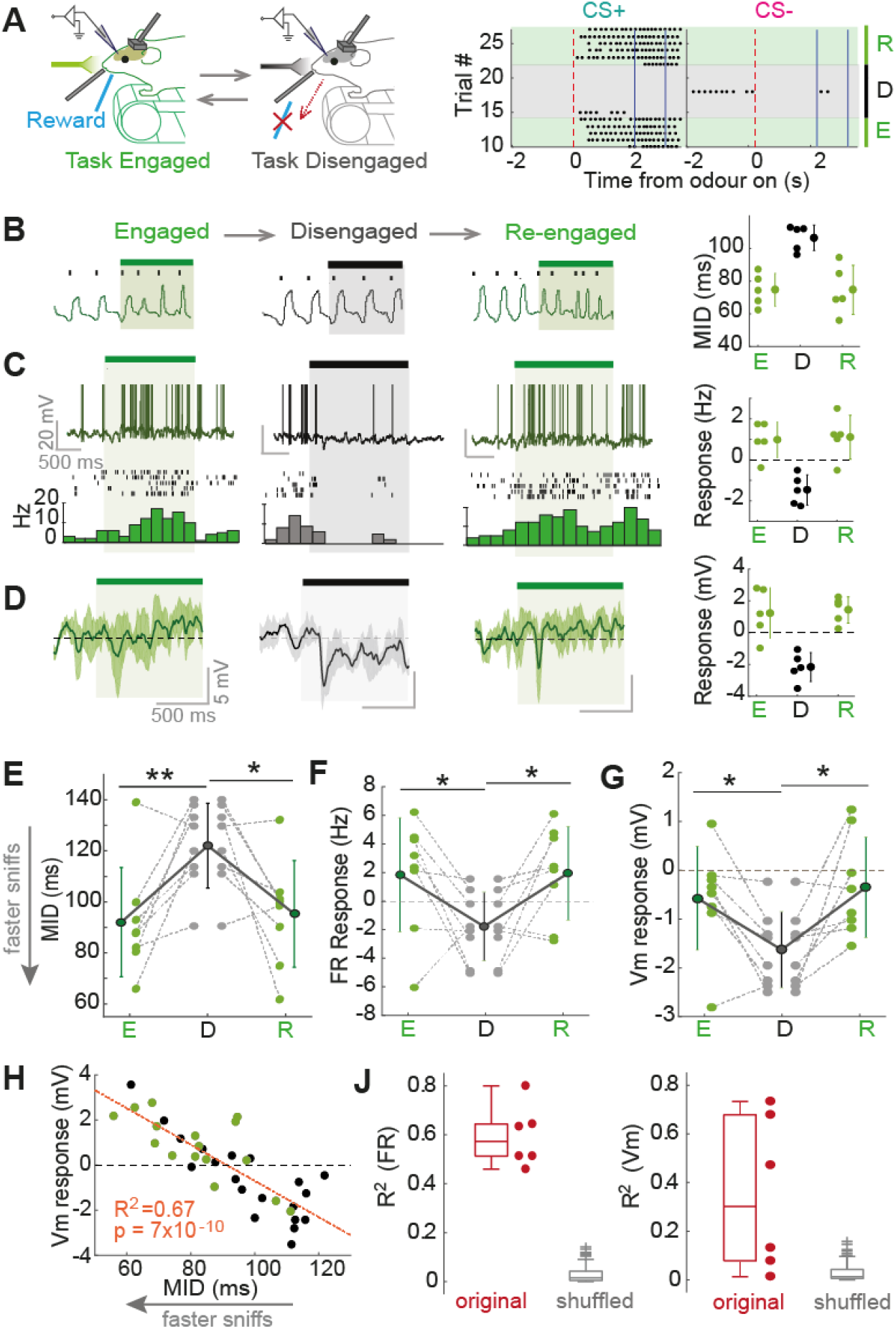
Active sampling and associated response changes are dynamically linked to task engagement. For all panels: green = task engaged, black = task disengaged. (**A**) Left shows experimental paradigm. Right shows lick raster across task switches for CS+ and CS− stimuli for an example mouse. (**B**) Example MID changes for one response across changes in task engagement (averaged over 500 ms, panels B-D correspond to same example). (**C**) Example FR response changes, (spikes partially clipped for display, averaged over 2 s). (**D**) Example V_m_ response changes (spike-subtracted, averaged over 500 ms). (**E**) For all 8 responses, changes in MID between task engagement, disengagement and re-engagement (asterisks denote result of paired T-test). (**F**) As for E, but for changes in 2 s FR responses. (**G**) As for E, but for changes in 500 ms V_m_ responses. (**H**) Scatter of MID versus V_m_ response across trials for an example cell-odor pair. (**J**) Left: boxplots to show corresponding R^2^ values (as for example in panel H) for all six FR responses showing significant changes across engagement shifts, alongside shuffle control. Right: as for left, but for V_m_ responses.

If active sampling determines positive response change as predicted from learning mice (Figure 3), we would expect positive changes to occur alongside the rapid sniffing strategy. We found that responses could change robustly and reversibly between task engagement, disengagement and re-engagement, with some examples showing dramatic and reversible switches between excitation and inhibition (Figure 5C-D and S8C). Consistent with the learning-related changes, positive changes always occurred alongside reduced MID (Figure 5E-G) and were again tightly linked to the sniff changes on a trial-by-trial basis (Figure 5H-J), consistent with the idea that changes in neural responses are directly driven by sniff strategy. Strikingly, response changes could occur within only a single trial upon recognition of task re-engagement (Figure S8D) emphasizing the dynamic nature with which changes in active sampling state influence neural responses.

### Odor response changes associated with active sampling are dependent on behavioral state

We next wanted to assess whether the response changes observed during active sampling require attention to an olfactory stimulus, or whether any similar change in sniffing would cause the same response change regardless of behavioral state.

MTC activity is strongly patterned by sensory input locked to the sniff cycle in anaesthetized mice, giving rise to sniff-coupling of membrane potential (Adrian, 1950; Cang and Isaacson, 2003; Fukunaga et al., 2012; Macrides and Chorover, 1972; Margrie and Schaefer, 2003). Similarly, in our awake animals, we found that membrane potential during odor stimulation showed widespread modulation by the sniff cycle, with a variety of sniff-V_m_ modulation amplitudes (Figure 1E). Thus, it is possible that changes in response occurring with rapid sniffing at least partially result from bottom-up changes in the sniff-locked input pattern from OSNs.

We thus assessed whether evoking changes in sniffing similar to those observed in behaving animals could directly elicit response changes even in the absence of olfactory behavior. We found that unexpected whisker stimulation briefly increased sniff rates in passive mice (Figure 6A), quantitatively reproducing (and even exceeding) the sniff changes seen during learning (Figure S9). When paired with odor delivery, this resulted in a variety of largely positive odor response changes (ΔV_m_=+0.65±0.82 mV, p=0.03, paired T-test, n=10; Figure 6B). If these are mediated by bottom-up effects on the sniff-locked input, we may expect the changes to correlate with the degree to which the response is sniff-coupled. Indeed, the response changes were strongly correlated with the amplitude of sniff-V_m_ modulation, such that highly sniff locked cells underwent the largest changes when sniffing was altered (Figure 6C). These response changes are unlikely to be due to changes in arousal or from somatosensory input, since they were similarly present in anaesthetized mice, where using a double tracheotomy the frequency of artificial sniffing (airflow through the nose) could be altered independent of free tracheal breathing (Figure 6D-E). Response changes in anaesthetized mice were also significantly correlated with sniff-V_m_ modulation amplitude (R^2^=0.71, p=0.006, n=9; Figure 6F). Thus, in absence of olfactory behavior, evoking sniff changes results in response changes which depend on the amount of sniff-locked input to the cell.

**Figure 6.**
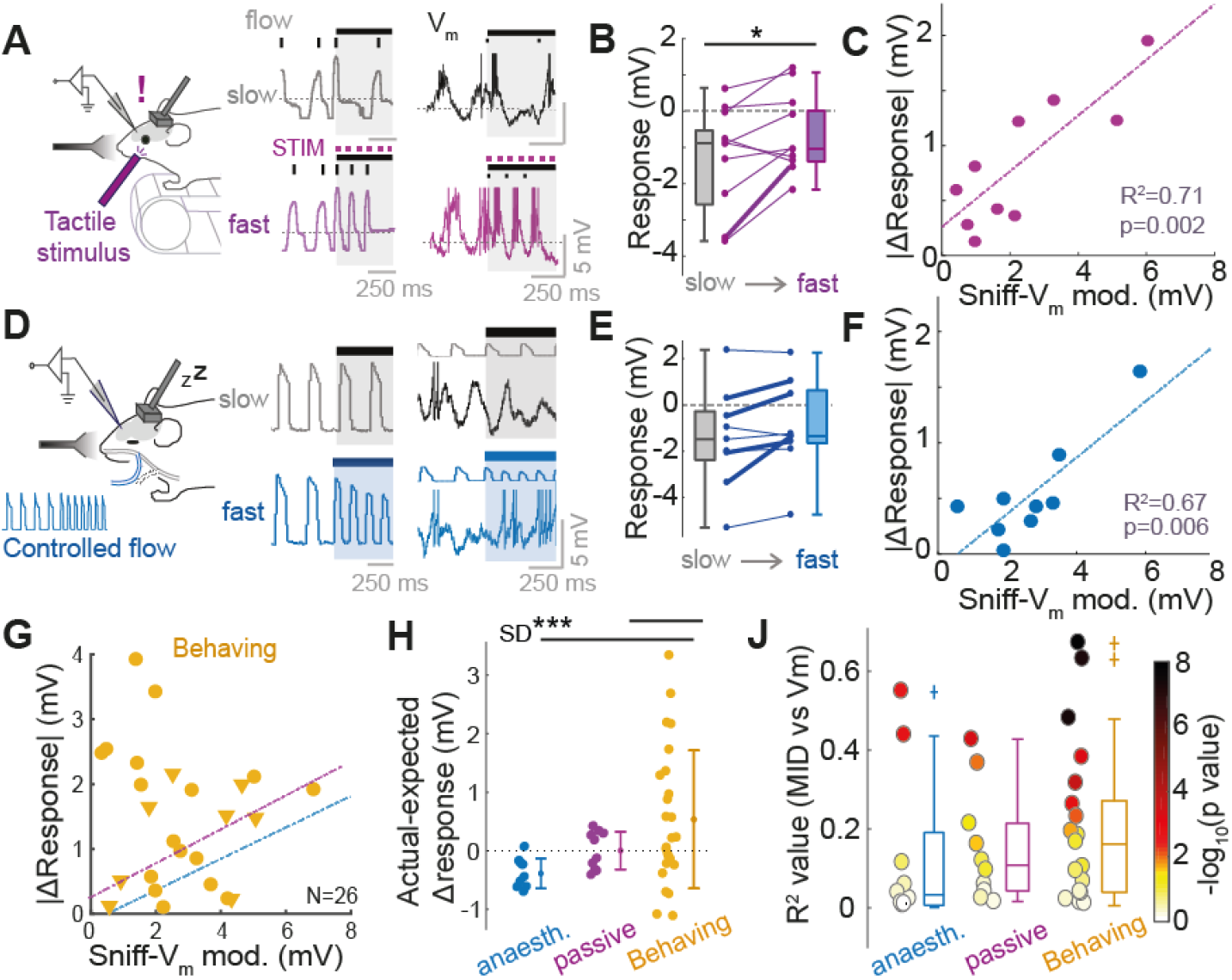
Response changes associated with active sampling are dependent on behavioral state. (**A**) Left: Experimental set up for tactile stimulation of passive mice. Example traces show nasal flow (middle) and Vm (right) for one cell: top = control trial (slow sniffing); bottom = tactile stimulus trial (fast sniffing). (**B**) Mean responses (first 500 ms of stimulus) averaged across 5 ‘slow’ and 5 ‘fast’ sniff trials for 10 cell-odor pairs (p<0.05, paired T-test). (**C**) Scatter of response change between slow and fast sniff trials versus odor sniff-V_m_ modulation amplitude during the odor. (**D**)-(**F**) as for the panels above, but for data from ‘simulated’ sniff changes in anaesthetized mice via a double tracheotomy procedure. (**G**) As for panels C and F, but for mice showing large sniff changes across learning (MID>20 ms) Correlations from C and F have been included for comparison. (**H**) Plot comparing deviation of response change from the linear regression model calculated from passive mice (linear fit in panel E), for anaesthetised, passive and behaving mice. (**J**) Comparion of R^2^ values between MID and V_m_ response calculated across trials for each cell odor pair undergoing large MID change in anaesthetised, passive and behaving mice.

We next wanted to assess whether this was the case in behaving mice. We thus pooled response changes from learning and task-engagement mice where MID underwent a change exceeding 20 ms (early-late or engaged-disengaged respectively). This gave us 26 cell-odor pairs in total. Plotting the absolute response change against the sniff-modulation amplitude resulted in a considerably different picture compared to passive and anaesthetized mice: there was no correlation between sniff-V_m_ modulation strength and response change magnitude (R^2^=0.02, p=0.6, n=26; Figure 6G). Using the linear model resulting from the correlation calculated in passive mice (ΔV_ex_=0.26\**T*+0.21 mV, where ΔV_ex_=expected V_m_ response change, and *T*= sniff-V_m_ modulation amplitude), we generated expected values for V_m_ response change based on the sniff-V_m_ modulation amplitude of each cell odor pair, and compared these to actual values for response change. On average, only response changes in behaving mice exceeded that expected based on their sniff-V_m_ modulation amplitude (mean actual-expected error=0.5±1.2 mV, p=0.03, paired t-test), and there was significantly more variance in the prediction error for response changes in behaving animals (actual-expected SD=1.2 mV) relative to passive (SD=0.3, p=3×10^−4^) and anaesthetized mice (SD=0.26, p=1×10^−4^; Figure 6H).

While this suggests that sniff-evoked response changes in behaving mice exceed those expected based purely on sniff-locked feed-forward input, this does not mean that such response changes are any less linked to the sampling behavior of the animal. When comparing R^2^ values for the correlation between MID and V_m_ response across trials for each cell-odor pair, we found no significant differences in the distributions between anaesthetized, passive and behaving mice (Figure 6J), and the latter if anything displayed larger R^2^ values (behaving: median=0.16, IQR=0.03-0.27; passive: median=0.10, IQR=0.03-0.21; anaesthetised: median=0.03, IQR=0.01-0.19; p>0.05, ranksum) and more frequent significant relationships (behaving: 46%; passive: 40%; anaesthetised: 20%; p<0.05 linear regression).

Thus, sniff changes in all behavioral states will evoke response changes to a degree, but in the behaving, actively sampling animal, these changes exceed those expected based only on the feed-forward input to the cell. This likely indicates a state-dependent top-down component underlying response changes during active sampling.

### Effect of fast sniffing in absence of odor depends on feed-forward input in learning and passive mice

A previous study in the visual system has shown that modulation of visual responses happens temporally locked to saccade generation (Han et al., 2009). Since activity even in absence of odor is widely modulated by the sniff cycle (Cang and Isaacson, 2003; Fukunaga et al., 2012; Macrides and Chorover, 1972; Figure S4F), and sniff changes evoke activity changes in all behavioural states given that they are highly sniff-coupled (Figure 6), this made it likely that sniff changes even in absence of odor would cause activity changes. We wanted to test whether the enhancement of response change during active sampling (Figure 6G-H) occurred only during the odor stimulus, or whether there is a generally increased sensitivity to sniff changes during behavior that extends outside the stimulus sampling period.

To examine this, we made use of spontaneous bouts of rapid (>5 Hz) sniffing that occur in awake mice during the inter-trial interval – i.e. in absence of odor (Figure 7A). Consistent with previous imaging data (Kato et al., 2013), it was clear that in certain cells, overt depolarising and hyperpolarising changes in activity would occur coinciding with such rapid sniff bouts (Figure 7A). Quantifying the change in mean membrane potential evoked by fast sniffing across 26 MTCs revealed almost two thirds significantly changed their mean potential during fast sniffing, with 7 depolarizing and 9 hyperpolarizing (p<0.05, bootstrapping – see methods, Figure 7A). Thus, sniff changes evoke response changes even in absence of odor. To test how these depended on bottom-up sniff-locked input, we again compared the magnitude of the response changes to their sniff-V_m_ modulation amplitudes. This resulted in a robust correlation (R^2^=0.46, p=0.001, n=26; Figure 7B), indicating that these changes are again likely the result of changes in feed-forward input.

**Figure 7.**
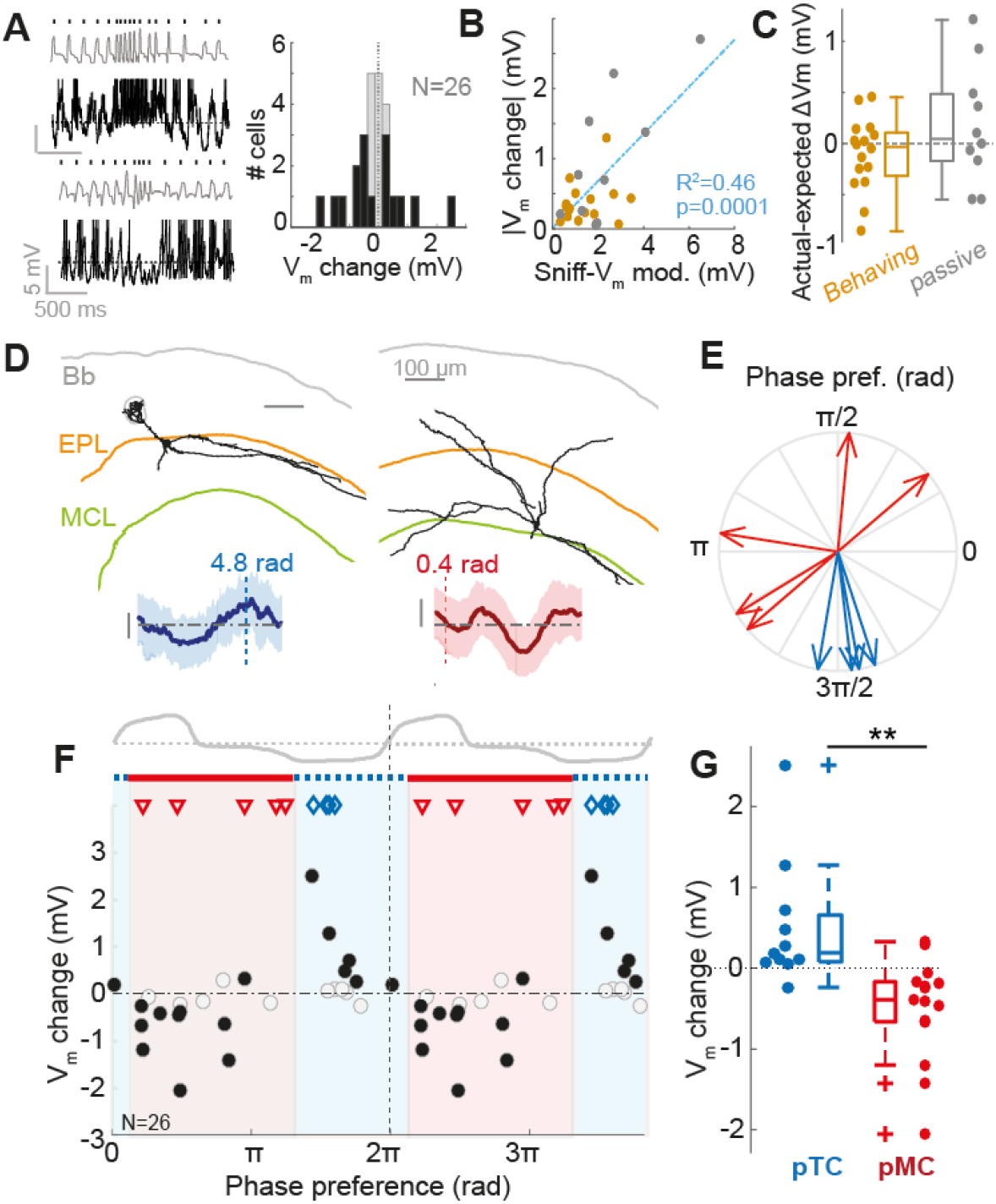
Response changes in absence of odor are dependent on sniff-locked input. (**A**) Right: awake mice will sometimes make spontaneous rapid sniff bouts in absence of odor during the inter-trial interval. Example traces show such sniff bouts, and coincident V_m_ traces showing overt activity changes. Left: Histogram to show distribution of V_m_ changes during spontaneous rapid sniffs (>5 Hz) for 26 MTCs in which there were >20 fast sniffs. (**B**) Correlation between absolute sniff change between slow and fast sniffs, and amplitude of baseline theta modulation. Grey dots show data from passive mice (n=10), gold dots show data from behaving mice (n=16). (**C**) Comparison of errors (actual-expected) when calculating expected V_m_ change based on the sniff-V_m_ modulation amplitude of the baseline membrane potential for passive (grey), and behaving (gold) cell-odor pairs. (**D**) Morphologies of a reconstructed TC (left) and MC (right), with mean membrane potential as a function of phase shown below (shaded area=SD), with their respective phase preferences. Bb=brain border; EPL=external plexiform layer; MCL=mitral cell layer. (**E**) Phase plot to show preferences of 5 reconstructed MCs (red) and 4 reconstructed TCs (blue). (**F**) V_m_ change between fast and slow sniffing (fast-slow) as a function of the phase preference of the cell. Red shaded region corresponding to inhalation and subsequent pause shows the phases which best encompass hyperpolarising cells, thought to be MCs, and blue region best encompasses depolarising cells, thought to be TCs. Symbols show phase preferences of morphologically recovered cells: red triangles=MCs; blue diamonds=TCs. Black filled dots show significant V_m_ changes. (**G**) Comparison of V_m_ change due to fast sniffing for putative TCs and MCs defined by the phase boundaries shown in panel F.

To test any differences in sensitivity to sniff change caused by behavioral state, we split the data into those from behaving mice (n=16) and those from passive mice (n=10). Comparing the actual V_m_ change to the expected V_m_ change (as calculated using the linear model generated from the linear regression, ΔV_ex_=0.31\**T*+0.01 mV, where ΔV_ex_=expected absolute V_m_ change, and *T*= sniff-V_m_ modulation amplitude), showed that the difference between expected and actual V_m_ change did not significantly differ from zero for either passive (mean actual-expected=0.17±0.57 mV, p=0.37, paired t-test, n=10) or behaving cell-odor pairs (mean actual-expected=−0.11±0.36 mV, p=0.25, paired t-test, n=16; Figure 7C), and did not significantly differ between passive and behaving datasets (p=0.14, unpaired t-test; p=0.1, Bartlett test). Altogether this indicates that enhanced response change during rapid sniffing in a behaving animal is only true during the odor sampling period.

Since cells could depolarise or hyperpolarise during fast sniffing, we sought to determine whether the sign of response change was also predictable from sniff-locking properties. Evidence from anaesthetized mice suggests that MCs are driven by feed-forward inhibition and lock to inhalation, while TCs are driven by feed-forward excitation and lock to exhalation (Fukunaga et al., 2012). To test this in awake mice, we recovered 9 morphologies of MTCs (e.g. Figure 7D), and identified them as MCs (n=5) or TCs (n=4). Congruent with the previous data, the two cell types had subthreshold membrane potential which locked to different phases of the sniff cycle: morphologically-confirmed MCs locked to inhalation, while TCs locked to exhalation (Figure 7E). We next examined the relationship between phase preference and the effect of fast sniffing across the full sample of cells. The sign of the change in activity during fast sniffs was strongly related to the phase coupling of the cell to the sniff cycle (Figure 7F), with inhalation-locked cells hyperpolarising and exhalation-locked cells depolarising. We calculated the phase boundaries for best separation of hyperpolarising and depolarising cells (as drawn in Figure 7F; see methods), and the phase preferences of morphologically identified MCs and TCs conformed to these boundaries (Figure 7F, red triangles and blue diamonds). Cells within the inhalation boundaries (0.39-4.11 rad; putative MCs) showed significantly more hyperpolarising effects of fast sniffing than those within the exhalation boundaries (4.11-0.39 rad; putative TCs) (putative MC, median ΔV_m_=−0.39 mV, IQR=−0.66 to −0.17 mV, n=16; putative TC median ΔV_m_= 0.19, IQR=0.08-0.66, n=11; p=9×10^−4^, Ranksum; Figure 7G).

Thus, in absence of odor, the effect of fast sniffing on response is predicted by the sniff-driven input of the cell regardless of behavioural state, and the sign of response allows identification of putative MCs and TCs.

### Tufted cells show more highly correlated changes than mitral cells

Since our data suggests involvement of extrabulbar circuits in shaping responses during active sampling, and previous work has suggested that both learning and neuromodulatory activity may have divergent effects on MC responses compared to TC responses (Kapoor et al., 2016; Yamada et al., 2017), we wanted to compare the response changes across learning for the two groups of cells. To this end we used the phase preference boundaries found earlier (Figure 7F) to designate putative mitral (pMC) and tufted cell (pTC) phenotype. Consistent with the idea that these boundaries can separate TCs and MCs, mean firing rate responses to odors in pTCs showed a significant tendency toward strong excitation compared to pMCs (Figure S10), as has previously been demonstrated (Nagayama et al., 2004).

The distribution of early subthreshold responses (prior to learning) for pMCs and pTCs did not significantly differ (pTCs: −1.1±1.9 mV, n=16; pMCs: −1.8±2 mV n=26; p=0.26, unpaired t-test; Figure 8A), however pTCs showed significantly more positive responses compared to pMCs in late responses after learning was complete (pTCs: median=0.3 mV, IQR=−1.3-1.1 mV, n=16; pMCs: median=−2.1 mV, IQR=−3.2-0.5 mV, n=26; p=0.01, Ranksum), consistent with previous findings that TCs show more excitatory responses and receive less lateral inhibition than MCs (Christie et al., 2001; Nagayama et al., 2004). Comparing response changes across learning for putative MCs and TCs, we found that the two groups did not significantly differ in terms of mean or variance of response change (pTCs: 0.64±1.7 mV; pMCs: −0.14±1.4 mV; p=0.1, unpaired t-test; p=0.46, Bartlett test; Figure 8B). Comparing the R^2^ values for the correlations between inhalation duration and V_m_ response across trials also indicated that in general, pMCs and pTCs do not show differing effects of sniffing on responses (pTCs: median R^2^=0.09, IQR=0.01-0.29; pMCs: median R^2^= 0.06, IQR=0.03− 0.18; p=0.88, Ranksum; p=0.35, Brown-Forsythe test; Figure 8C).

**Figure 8.**
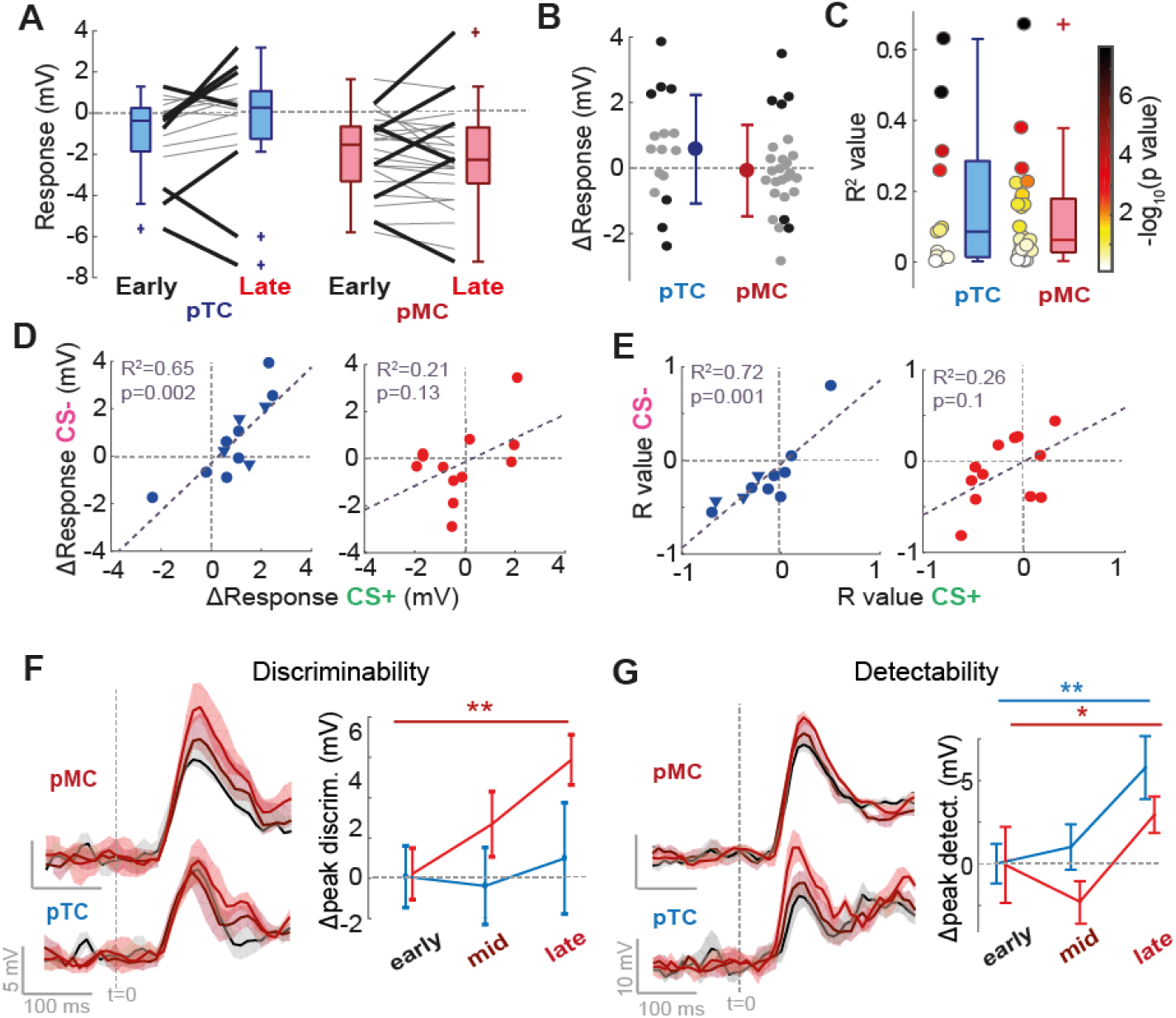
Tufted cells show more highly correlated changes than mitral cells. (**A**) Plot of early and late membrane potential responses (first 500 ms) across learning for pTCs (left; n=16 cell-odor pairs) and pMCs (right; n=26 cell-odor pairs) separately. Lick black lines show significant changes. (**B**) Comparison of response changes (late-early) for pTCs and pMCs cell-odor pairs. Black dots show significant changes (p<0.01). (**C**) Comparison of R^2^ values between MID and V_m_ response across trials for pTC and pMC cell-odor pairs. Color shows p-value of the correlation (−log10). (**D**) Scatter of response change for CS+ vs response change for CS− for pTCs (left) and pMCs (right) independently. Triangles show data from task-engaged/disengaged recordings (as in Figure 5), where response change is calculated as engaged-disengaged response, while circles are from learning data (response change=late-early response). (**E**) As for G, but for the R value between MID and V_m_ response across trials for each cell odor pair. (**F**) Left: Euclidean distances for the discriminability between CS+ and CS− during early (black), mid (maroon) and late (red) trials, for pTCs (bottom) and pMCs (top) independently. Right: Left: plot of average peak discriminability in the first 170 ms of the stimulus for early, mid and late trials for pTCs (blue) and pMCs (red) Plot shows mean, and error bars show SD across 5 trial subsets. (**G**) Right: Euclidean distances (as in Figure 4) for the detectability between CS+ and CS− during early (black), mid (maroon) and late (red) trials, for pTCs (bottom) and pMCs (top) independently. Left: plot of average peak detectability in the first 170 ms of the stimulus for early, mid and late trials for pTCs (blue) and pMCs (red) Plot shows mean, and errorbars show SD across 5 trial subsets.

We next compared the response changes for CS+ and CS− stimuli across learning for pMCs and pTCs individually. For tufted cells, response changes for the two stimuli were highly significantly correlated (R^2^=0.65, p=0.002, n=12 cells), whereas this was not the case for pMCs (R^2^=0.21, p=0.13, n=13 cells; Figure 8D). The same difference was seen when looking at the R values between MID and V_m_ response across trials: pTCs showed highly correlated R values between CS+ and CS− stimuli (R^2^=0.72, p=0.001, n=11), while pMCs did not (R^2^=0.26, p=0.1, n=12; Figure 8E).

Since response changes were overall less correlated between CS+ and CS− for MCs, we wanted to compare the change in discriminability of the responses across learning for pMCs compared to pTCs. Using the Euclidean distance between population response vectors for CS+ and CS− stimuli, we found that pTCs did not show a significant change in peak discriminability across learning (mean peak early=9.4±1.6; late=10.4±2.6 mV, p=0.41 unpaired t-test, n=5), however pMCs did show a significant increase in peak discriminability (mean peak early=10.1±0.4; late=13.1±1.9 mV, p=0.01 unpaired t-test, n=5; Figure 8F). Both cell types however significantly contributed to increased detectability of the stimulus across learning, though this was more pronounced for TCs rather than MCs (TCs: peak early= 15.9±1.2 mV, peak late=21.5±1.8 mV, p=0.001, unpaired t-test; MCs: peak early=31.0±2.2 mV, peak late= 33.9±1.1 mV, p=0.01, unpaired t-test; Figure 8G).

Thus, while response changes across learning were generally quite similar for MCs and TCs, TCs showed more highly correlated changes while the less correlated changes in MCs appear to enhance discriminability of the stimuli.

## Discussion

Active sampling behavior is a fundamental feature of sensory information acquisition. Theoretical and psychophysical evidence has driven hypotheses that active sampling strategies during behavior may be used to optimize sensory information flow (Ahissar and Assa, 2016; Laing, 1983; Yang et al., 2016). Here using whole-cell recordings in awake mice, we found a number of differences in subthreshold responses between passive and learning mice (Figure 1), with variance in responses developing across the rapid learning episode (Figure 2). In parallel, we found that active sniffing develops across learning in motivated mice (Figure 3), which corresponds to changes in odor response (Figure 4 and 5), ultimately serving to improve odor representation. Moreover, we show that this cannot be predicted from simple feed-forward mechanisms (Figure 6), a feature which only holds true during odor sampling (Figure 7), and occurs in a cell-type specific manner (Figure 8). Thus, we provide new evidence for coordinated modulation of early sensory processing during active sampling epochs, which serves to enhance early odor representation.

Rodents alter their sniffing pattern in all kinds of contexts (Wachowiak, 2011), both in absence of odor (Bramble and Carrier, 1983; Ikemoto and Panksepp, 1994; Wesson et al., 2008), as well as during odor sampling in behavioral tasks (Kepecs et al., 2007; Roland et al., 2016; Wesson et al., 2009; Youngentob et al., 1987). We show that a portion of the variance in odor sampling strategy can be explained by motivational state (Figure 3J-K). Thus, active sampling strategies are highly context dependent. Sniff changes will have an overt effect on highly sniff locked cells in absence of olfactory behavior dependent on their feed-forward input (Figure 6C and F), and an even more profound effect on a wider range of cells if the animal is engaged in odor-directed active sampling (Figure 6G-H). As such, the precise effect of sniff changes on mitral/tufted cell activity is itself dependent on behavioral context. Changes in sniffing strategy could therefore provide a common mechanistic basis for a number of different contextual modulations described in OB activity (Beshel et al., 2007; Chu et al., 2016; Di Prisco and Freeman, 1985; Freeman and Schneider, 1982; Fuentes et al., 2008; Kay and Laurent, 1999; Pager et al., 1972; Rinberg et al., 2006; Doucette and Restrepo 2008). However, we note that some variance in response cannot be explained by active sampling, such as the increases in inhibition during learning we note here (Figure 4E and F). The enhancement of odor representation seen during active sampling could explain the improvement in discrimination performance previously reported for mice displaying active sniffing strategies (Kepecs et al., 2007), as well as the faster reaction times noted here (Figure 3L).

Odor responses during active stimulus sampling are enhanced compared to changes seen during sniff changes in absence of olfactory behavior (Figure 6). This suggests the involvement of top-down centers that serve to coordinate sensory processing at the periphery with the active sampling state of the animal (Wachowiak et al., 2011). Congruently, several neuromodulatory centers which project to the OB interact with respiratory control centers in the brainstem, including serotonergic fibers and the noradrenergic locus coeruleus (Dugué and Mainen, 2009; Yackle et al., 2017). We find a cell-type specificity in the effect of active sampling on response changes congruent with recent imaging across learning (Yamada et al., 2017), and neuromodulatory centers have recently been shown to have divergent effects on MCs and TCs – with serotonin having more heterogeneous effects on MCs and TCs (Kapoor et al., 2016). As such, neuromodulators are a prime candidate to coordinate OB state with active sampling behavior. In the whisker system, cholinergic afferents in the barrel cortex are known to be active during spontaneous whisking and mediate changes in physiology (Eggermann et al., 2014), while *in vivo* activation of these afferents boosts sensory input to the OB (Bendahmane et al., 2016). Future investigation will be required to address which centers are activated during active sampling, alongside their targets within the olfactory bulb circuit.

Complex orchestration of active sampling is similarly present in other sensory systems; whisking shows modulations during exploratory behavior (Mitchinson et al., 2007), and eye movement varies between tasks and individuals (Hayhoe and Ballard, 2014; Rayner et al., 2007) with both behaviours affecting sensory cortical activity (Crochet and Petersen, 2006; McFarland et al., 2015). Whether and how directed adjustments to such active sampling during behavior might also improve early sensory representations in these other modalities remains to be seen.

In conclusion, early sensory activity in the olfactory bulb is modulated by dynamic adjustments in the closed-loop pathway that coordinates active sniffing (Ahissar and Assa, 2016), yielding enhanced sensory representation during olfactory behavior.

## Author contributions

A.T.S., R.J., I.F. and M.K. designed all experiments, R.J. performed all experiments, and analysed all data with help from A.T.S., I.F. and M.K. R.J. and A.T.S. wrote the article with contributions from I.F. and M.K. The authors declare no competing financial interests.

## Acknowledgements

We thank Martyn Stopps and Nicholas Burczyk for assistance with custom made equipment, Mostafa Nashaat and Edward Bracey for initial support with behavioral training, Christoph Schmidt-Hieber for advice on whole-cell recording *in vivo*, and Roma Shusterman, Alexander Fleischmann, Kevin Franks, Mahesh Karnani, Denis Burdakov, Michael Hausser, Tim Gollisch, Tobias Ackels and Ede Rancz for helpful comments on the manuscript. This work was supported by the Francis Crick Institute which receives its core funding from Cancer Research UK (FC001153), the UK Medical Research Council (FC001153), and the Wellcome Trust (FC001153); by the UK Medical Research Council (grant reference MC_UP_1202/5); by the DFG (SPP 1392); and a Boehringer Ingelheim Fonds PhD fellowship to RJ. AS is a Wellcome Trust Investigator (110174/Z/15/Z).

## Methods

All animal experiments were approved by the local ethics panel of the Francis Crick Institute (previously National Institute of Medical Research) and UK Home Office under the Animals (Scientific Procedures) Act 1986. All mice used were C57BL/6 Jax males aged between 5 and 8 weeks obtained by in house breeding. All chemicals were obtained from Sigma-Aldrich (Missouri, USA).

### Head-fixation

For surgical procedures, strict sterile technique was adhered to. Mice were anaesthetized with isoflurane in 95% oxygen (5% for induction, 1.5-3% for maintenance), and received general analgesia (Carprofen, 5mg/kg s.c.) as well as local analgesia around the dorsal surface of the head (Levobupivicaine or Mepivicaine, 0.5% s.c.). A custom-made stainless steel headplate was affixed to the intraparietal and parietal skull plates with a combination of cyanoacrylate and dental cement, while a recording chamber was constructed upon the bone overlying the right olfactory bulb using a plastic ring and dental cement. The chamber was filled with silicone (Quik-Cast - World Precision Instruments, Florida, USA) and sealed during the recovery and training periods prior to recordings. After 48 hours recovery, mice going on to passive experiments were head-fixed under very light isoflurane anesthezia (identical to the trained mice, see below) and allowed to awaken on a custom-made treadmill. Mice were allowed to accustom themselves to the treadmill in this initial 20 minute session, by the end of which mice showed no stress behavior and learned to walk and sit calmly on the treadmill. Mice going on to behavioral training underwent 2 days of additional water scheduling prior to head-fixation, and in the initial head-fixation session were additionally allowed access to abundant free rewards (diluted sweetened condensed milk) upon licking at the reward spout.

### Go/No-Go behavior

The day following head-fixation habituation, mice undergoing go/no-go training progressed to two more days of pre-training for acquisition of the go/no-go task. On the first day mice were presented only the CS+ odor and were trained to acquire the ‘go’ licking pattern following odor offset via a delay classical conditioning procedure. Note that no measure was in place to prevent or punish licking behavior during the odor stimulus, and some mice would additionally lick during the odor stimulus prior to the allotted response time after odor offset (termed ‘anticipatory licking’). Following successful learning of this lick pattern, the next day mice were presented both the CS+ and CS− on a pseudorandom basis. Mice had to learn to respond to these odors differentially, learning to inhibit responses (‘no-go’ behavior) for the CS− to avoid a 5 s addition to the ITI. Only when mice had successfully demonstrated learning of this task (two consecutive 10-trial blocks of at least 80% correct responses) they were moved on to whole-cell recording procedures the next day. After successful acquisition of a recording, mice were presented a novel pair of odor stimuli assigned each to CS+ or CS−, and had to learn the go/no-go behavior for these new stimuli. Criterion within a recording was considered one block of at least 80% performance. Learning of the task with the second pair of stimuli was always far more rapid than for the original acquisition (Figure S2B), well within whole-cell recording timescale in awake mice. For mice undergoing the task engagement/disengagement paradigm, acquisition of the task occurred prior to recording such that criterion performance was already achieved from the start of the recording. After 20-30 trials, the water port was manually moved away to disengage the task. Mice would continue to attempt to lick (as detected by infrared beam) for a variable number of trials before ‘giving up’ (i.e. 5 consecutive ‘miss’ trials), after which the port was returned. Often a free reward was used as a salient stimulus to the mouse that the task was re-engaged.

### Odor delivery

Odor stimuli were delivered using a custom-made airflow dilution olfactometer with electronic dilution control. All odor stimuli were calibrated using a mini photoionization detector (miniPID, Aurora Scientific, Ontario, Canada) to form square-pulses of 1% concentration (relative to maximum recorded vapor-pressure in air, Figure S1). Odor stimuli used for initial go/no-go training purposes consisted of peppermint oil and almond oil - components that were not present in the odor mixtures later presented in recordings. For stimuli during whole-cell recordings, 2 were randomly selected from 4 potential odor mixtures (Figure S1), and for behaving mice randomly assigned to CS+ or CS−. Odor mixtures were comprised of 4 to 6 monomolecular odorants selected for their reported ability to activate dorsal glomeruli (Takahashi et al., 2004), grouped according to similarity of vapor pressure, and added to the mixture in an undiluted quantity inversely proportional to their relative vapor pressures (Figure S1). Odors were presented with a minimum inter-trial interval of 10 s. To minimize contamination, a high flow clean air stream was passed through the olfactometer manifolds during this time. Constant air-flow going to the animal was achieved using a final valve, minimizing any tactile component accompanying the odor stimulus.

### Whole-cell recordings

Animals were again anaesthetized under isoflurane as before, and recording chambers were re-opened. A 1-2 mm craniotomy and durectomy was made over the right olfactory bulb. The craniotomy was then covered with a 0.5-1 mm layer of 4% low melting-point agar, which greatly contributed to the stability of recordings. This layer was removed and re-applied after every descent of a recording micropipette. The recording chamber was then filled with cortex buffer (125 mM NaCl, 5 mM KCl, 10 mM HEPES, 2 mM MgSO_4_, 2 mM CaCl_2_, 10 mM glucose), and the mice were transitioned to head-fixation and allowed 30 minutes to recover from anesthezia. After this time, behaving animals would demonstrate retention of go/no-go behavior acquired the day previously prior to attempt for a recording. Micropipettes were prepared with a resistance of 5-8 MΩ from borosilicate glass (Hilgenberg, Malsfeld, Germany) capillaries, and filled with intracellular solution (130 mM KMeSO4, 10 mM HEPES, 7 mM KCl, 2 mM ATP-Na, 2 mM ATP-Mg, 0.5 mM GTP, 0.05 mM EGTA, and in some cases 10 mM biocytin). Signals were amplified using an Axoclamp 2B amplifier (Molecular devices – West Berkshire, UK) and digitized by a Micro 1401 (Cambridge Electronic Design – Cambridge, UK) at 25 kHz. Drift in membrane potential, corrected for by spike thresholds, between the start and end of recordings was 0.9±1 mV, with an average duration of 14±4 minutes, and access resistance of 36±19 MΩ.

### Sniff measurement

Sniffing behavior was recorded either with a pressure sensor or flow sensor (Sensortechnics – Rugby, UK), externally located in close proximity to the left naris (contralateral to recording side). The precise orientation relative to the nostril was manually optimized prior to each recording in order to acquire the full sniff waveform in spite of any movement of the naris.

### Double tracheotomy

Two mice were anaesthetized with ‘sleep-mix’ (0.05 mg/kg Fentanyl, 5 mg/kg Midazolam, 0.5 mg/kg Medetomidine), and both local and general analgesia applied as above for head-fixation. After the head-plate surgery, a double tracheotomy was performed by exposing the trachea and inserting two catheters, one directed to the lungs through which the mouse could freely breathe, and the other directed to the nasal passages through which flow was controlled. To mimic sniffing, a peristaltic pump (Ismatec, Wertheim, Germany) was used to generate flow inward through the nares, with a flow controller to buffer out fluctuations and the periodic opening of a 3-way valve used to simulate regular inhalations, either at 3.3 Hz (100 ms opening times), or 6.6 Hz (50 ms opening times).

### Neuronal numbers

Altogether we report here recordings from 66 mitral and tufted cells. We report data from 42 cell-odor pairs from behaving animals over the timescale of learning (21 cells from 20 animals), 46 cell-odor pairs from passively exposed animals (23 cells from 20 animals), 8 cell-odor pairs from animals undergoing the task engagement/disengagement paradigm (4 cells from 4 animals), 10 cell-odor pairs from passive mice undergoing the unexpected puff experiment (9 cells from 9 animals), and 9 cells from two anaesthetized mice with a double tracheotomy. None of these cohorts are overlapping. Of the cells from mice across learning, 2 were excluded from any sniff analysis due to poor sniff signals (resulting in 38 cell-odor pairs, 20 accompanied by small (<20 ms) sniff changes, 18 by large sniff changes), and 2 were excluded similarly from the passively exposed dataset (42 cell-odor pairs).

### Data analysis

All data was pre-processed in Spike2 version 7.1 (Cambridge Electronic Design – Cambridge, UK) and analyzed in Matlab 2015b (Mathworks - Massachusetts, USA) and R using custom scripts and functions.

### Statistics

In all cases, 5-95% confidence intervals were used to determine significance unless otherwise stated. In all figures, a single asterisk denotes p<0.05, double asterisk denoted p<0.01 and a triple asterisk denotes p<0.001. Where these are preceded by ‘SD’, the p-value refers to the variances rather than the averages of the datasets. Means and error bars showing a single standard deviation either side are used in all cases for normally distributed data of equal variance. Two-sided Student’s t-tests were used for comparison of means and Bartlett tests used to compare variances, unless otherwise stated. Boxplots are used to represent any other data (data comparisons of unequal variance, or non-normally distributed data), where median is plotted as a line within a box formed from 25^th^ (q1) and 75^th^ (q3) percentile. Points are drawn as outliers if they are larger than q3 + 1.5 × (q3 − q1) or smaller than q1 − 1.5 × (q3 − q1). For such data, Ranksum tests were used to compare the medians, and Browne-Forsythe tests used to compare variance, unless otherwise stated. To determine points of significant difference between cumulative histograms, a bootstrapping method was used. Firstly, data underlying the two histograms would be shuffled between datasets, and cumulative histograms would be calculated from these shuffled sets. The difference at each point between the two histograms would then be calculated. This was repeated 10,000 times, and the differences between the real cumulative histograms would then be compared to the shuffled distribution at each point. An arrow was drawn on the points at which the actual difference exceeded the 99th percentile of the shuffled distribution.

### Sniffing analysis

To extract inhalation durations, firstly inhalation peaks were detected as any peak above a certain threshold set according to the amplitude of the signal. Inhalation onset was set at the nearest point pre-peak that the flow trace crossed zero, while inhalation offset was set at the nearest point post-peak that the flow trace crossed zero. The distance between these points was taken as the inhalation duration. The mean inhalation duration for the first 500ms of each odor presentation was calculated from the duration of all complete inhalations within that time period.

### Principal cell identification

Mitral and tufted cells were distinguished from interneurons as previously (Kollo et al., 2014). The current data set was pooled with the entire data set of neurons recorded in the OB of awake mice acquired previously (Kollo et al., 2014), and independent component analysis was performed on the AHP waveform (2 to 25 ms from spike onset) to reveal three independent components, upon which hierarchical cluster analysis was used to band the cells into two groups, ‘principal’ and ‘other’. Based on cell morphologies from the previous data set, and an additional 12 acquired in the current data set, 100% of the 22 morphologies obtained from the ‘principal’ group were confirmed as mitral/tufted cells, while 86 % of the 11 morphologies from the ‘other’ group were confirmed interneurons. Morphologies from the current data set were acquired as previously (Fukunaga et al., 2012; Kollo et al., 2014): mice were perfused following recordings with cold phosphate-buffered saline, followed by 4% (wt/vol) paraformaldehyde solution in phosphate-buffered saline. Fixed olfactory bulbs were embedded in porcine gelatin (10% wt/vol), before being fixed overnight in 4% paraformaldehyde. The OBs were then cut into 150 μm slices with a vibratome (Thermo Scientific – Massachusetts, USA) and stained with avidin-biotinylated peroxidase (ABC kit − Vector Labs, California, USA) and the DAB reaction. Biocytin-stained cells were traced using a Neurolucida system (MBF Bioscience, Vermont, USA). Principal cells were identified via soma size, cell body location with respect to the mitral cell layer, an apical dendrite reaching the glomerular layer and lateral dendrites branching in the external plexiform layer. MCs were distinguished from TCs based on proximity to the mitral cell layer.

### Odor responses and changes

For all analyses, the first presentation of each odor was excluded due to the elicitation of high frequency sniffing by the novel odorant, which rapidly decayed by the second presentation (Wesson et al., 2008). **General response calculations**: All traces were aligned to first inhalation onset following final valve opening. For V_m_ response calculations, spike waveforms, including the AHP, were subtracted from the V_m_ trace (−5 to 20 ms after spike peak). Responses for each trial were calculated as the mean V_m_ within the first 500 ms post odor onset, normalized to the baseline membrane potential in the 2 s prior to odor onset. FR responses were calculated as the mean number of spikes per 0.25 s time bin in the first 500 ms post odor onset, normalized to that calculated for 2 s prior to odor onset. Significant responses were determined for both V_m_ and FR using a paired t-test to compare baseline and odor-evoked activity for all trials. **For response changes across learning**: Significant changes between early and late trials for each odor response were identified by comparing the five ‘early trials’ in block 1 (stimulus presentation #2 to 6), with the 5 last presentations of the stimulus (‘late trials’). Significant change was determined using an unpaired t-test, p<0.05. **To determine onset of response change**: For each response, the mean V_m_ response waveform calculated for early trials was subtracted from that calculated from late trials, to generate a response change waveform at each time-point from odor onset. This was then normalised by the standard deviation of this resulting waveform during the baseline period 2 s prior to odor onset. Response change onset was detected where the response change magnitude first exceeded 2 standard deviations and remained there for at least 50 ms. **For task engagement/disengagement changes**: The first 500 ms of the stimulus was analyzed for V_m_ responses, and the full 2 s for FR responses. 5 trials of initial engagement were defined as the last 5 trials of each stimulus prior to physical port removal, disengagement trials were defined as 5 trials with at least 3 consecutive misses within the block, and re-engagement trials were based on the first 5 trials of the stimulus after the mouse initiates licking after port return.

### Detectability and discriminability analysis

For each response, five mean V_m_ response waveforms were generated from different sets of 3 early trials, 3 mid-point trials (mid-point trial ± 2 trials either side) and 3 late trials (aligned to first inhalation post odor onset). 3 early, 3 mid-point, and 3 late corresponding mean baseline waveforms were made by averaging inhalation-triggered waveforms from the ITI. Population response vectors were then constructed from these mean response waveforms for all cell-odor pairs recorded. At each time point relative to inhalation onset, the Euclidean distance was calculated between response and baseline vectors, and this was repeated five times for each baseline vector to gain a mean detectability over time, and a standard deviation. Minimum detection times were calculated as the first time post-inhalation where the mean detectability exceeded 2.5 × the SD of the baseline mean detectability, and remained so for at least 50 ms. The average baseline Euclidean distance 200 ms prior to odor onset was subtracted from the trace, normalizing the baseline to zero. Peaks of detectability were defined as the maximum detectability within the first 170 ms after odor onset. Discriminability was analyzed similarly, however the response vectors used to calculate the Euclidean distances were calculated between CS+ and CS− mean V_m_ response waveforms for the five sets of early, mid-point and late trials, i.e. the Euclidean distance was generated between population responses for CS+ and CS− separately.

### Sniff-V_m_ modulation amplitudes and preferences

The sniff-V_m_ modulation properties of each cell were calculated as previously (Fukunaga et al., 2012). **Baseline sniff-V_m_ modulation:** due to the high variability of sniff behavior in awake mice, analysis was restricted to sniff cycles between 0.25 and 0.3s in duration, where also the preceding sniff cycle was within this range. Mean V_m_ from the spike-subtracted V_m_ trace was taken as a function of sniff cycle phase for at least 150 sniffs, and this was plotted as Cartesian coordinates. The angle of the mean vector calculated by averaging these Cartesian coordinates was taken as the phase preference of the cell, while the difference between the mean V_m_ at the preferred phase, and the minimum value on the mean V_m_ waveform was taken as the amplitude of modulation. **Odor sniff-V_m_ modulation:** This was calculated as for baseline, but based on the first four sniffs post odor onset for the 10 trials of lowest sniff rates. As odor responses can have both tonic and sniff-modulated components, the phase-V_m_ trace for each sniff had to be normalized according to the linear vector connecting the V_m_ at the beginning and end of the sniff. To determine significance, a bootstrapping method was used: 100 ms segments of V_m_ data were randomly selected for each cell and connected to form a shuffled dataset. The phase analysis was then performed on these shuffled datasets, and a modulation amplitude calculated and this was repeated 100 times. Significant modulation was assigned when the actual modulation amplitude exceeded that of the 95^th^ percentile of shuffled data amplitudes.

### Putative mitral cell versus tufted cell identification

For each ITI, the mean V_m_ was calculated during sniffs of duration of <200 ms where also the preceding sniff was within this duration range (‘fast sniffs’). This mean V_m_ was then normalized by subtracting the mean V_m_ during sniffs of duration 0.25 and 0.3s within the same ITI to calculate the ‘fast-sniff evoked V_m_’. Only cells with at least 20 such ‘fast sniffs’ within the recording were considered for the analysis. To determine significance, a bootstrapping method was used: the mean V_m_ for all sniffs within a trial was randomly shuffled, and the shuffled data analyzed as before 100 times. The actual fast-sniff evoked V_m_ was then compared to the 5^th^ and 95^th^ percentiles of the shuffled distribution in order to assign significance.

We noted that, consistent with anaesthetized mice (Fukunaga et al., 2012), there was a bimodal distribution of phase preferences for the sniff cycle in baseline membrane potential, one within exhalation phase, and another within inhalation phase. We hypothesized that these may correspond to MC and TC phenotypes respectively, as reported previously for anaesthetized animals (Fukunaga et al., 2012). The putative assignment to MC or TC was confirmed morphologically for 8 cells (Figure 7F), with MC and TC distinction based largely on soma location relative to the mitral cell layer, as dendritic reconstruction was in many cases incomplete (Fukunaga et al., 2012).

### Unexpected tactile stimulus experiments in passive mice

In 10 passive mice, odors were presented as before, but this time with a random chance of an unexpected tactile stimulus to accompany the odor (25% chance) to evoke fast sniffing. Since the sniffing response to the tactile stimulus eventually habituated, for each response, the five trials with lowest MID were selected and compared to the five trials with highest MID. The difference in response for these sets of trials was then calculated for the first 500 ms of the stimulus as for learning mice.

### Reaction times

Reaction time calculations were based on 10 or more trials of 80% performance. **From lick behavior**: For each CS+ and CS−, lick probability was calculated in a moving time window of 100 ms, aligned to the first inhalation after final valve opening. The difference between the probability of licking for CS+ and CS− for each time window was calculated, and the leading edge of the first window at which this calculated difference significantly deviated from the values calculated from the 2 s window prior to odor onset was considered the reaction time (Figure S2C). **From sniff behavior**: Inhalation and exhalation duration values were calculated for CS+ and CS− as a function of sniff number from odor onset. These values were compared between those calculated for CS+ and CS− using a t-test, and the decision time was calculated based on the first inhalation or exhalation within the series to show a significant difference (Figure S2D). For 12/21 mice there was a significant difference between CS+ and CS− sniffing.

### Response onset analysis

For each response, the mean V_m_ response waveform calculated for early trials was subtracted from that calculated from late trials, to generate a response change waveform at each time-point from odor onset. This was then normalized by the standard deviation of this resulting waveform during the baseline period 2 s prior to odor onset. Response change onset was detected where the response change magnitude first exceeded 2 standard deviations and remained there for at least 50 ms. To determine the effect of sniff changes on response onset, only the first inhalation after odor onset was considered, since only response onsets ≤250 ms – within the first sniff cycle – were analyzed. For each response, trials were categorized into ‘slow’ (>90 ms inhalation duration) or ‘fast’ (<90 ms) sniff trials. The mean normalized V_m_ response waveform was averaged across these trials. Response onsets were calculated as before using these waveforms. Only cases where there were 5 or more trials in each category were analyzed. Cases where the mean 500 ms V_m_ response for either slow or fast sniffs was less than 0.5 mV in amplitude were also discarded. Response onsets from fast vs. slow trials were then compared across all responses for either behaving or passive mitral/tufted cells only to determine the effect of sniffing within each group. Response onsets were then compared between passive and behaving mitral/tufted cells for either slow or fast sniffs only, to determine any effect of behavioral state independent of sniff duration. To determine significant differences, a paired T-test was implemented for slow vs fast sniff groups within passive or behaving cohorts, or Ranksum tests were used when comparing between passive and behaving cohorts.

